# Implication of HisF from *Acinetobacter baumannii* in persistence during a pneumonia infection

**DOI:** 10.1101/534966

**Authors:** Marta Martínez-Guitián, Juan C. Vázquez-Ucha, Laura Álvarez-Fraga, Kelly Conde-Pérez, Juan A. Vallejo, Germán Bou, Margarita Poza, Alejandro Beceiro

## Abstract

The *hisF* gene from *A. baumannii* ATCC 17978 was found over-expressed during a murine pneumonia infection. A mutant strain lacking *hisF* showed its involvement in virulence during mice pneumonia as well as in host inflammatory response, where the product of HisF may act as negative regulator in the production of pro-inflammatory cytokines. This work evaluates the role of HisF in the *A. baumannii* pathogenesis and suggests its potential as a new target for antimicrobial therapies.

## TEXT

*Acinetobacter baumannii,* included by the WHO in a list of the most important antibiotic resistant pathogens (1, 2), shows a great capacity to persist in the hospital environments developing antimicrobial resistance. There is an urgent need of finding new therapeutic targets for the design of new strategies for fighting against this bacterium.

The *hisF* gene of *A. baumannii* is involved in purines and histidine biosynthesis. The *hisH* and *hisF* products shape the heterodimeric protein imidazole glycerol phosphate (IGP) synthase. This heterodimeric enzyme catalyzes the transformation of the intermediate N’-(5’- phosphoribosyl)-formimino-5-aminoimidazol-4-carboxamide ribonucleotide (PRFAR) into 5’- (5-aminoimidazole-4-carboxamide) ribonucleotide (AICAR) and imidazole glycerol phosphate (ImGP), which are further used in purine and histidine biosynthesis, respectively(3-5) (Figure 1).

**Figure 1.**
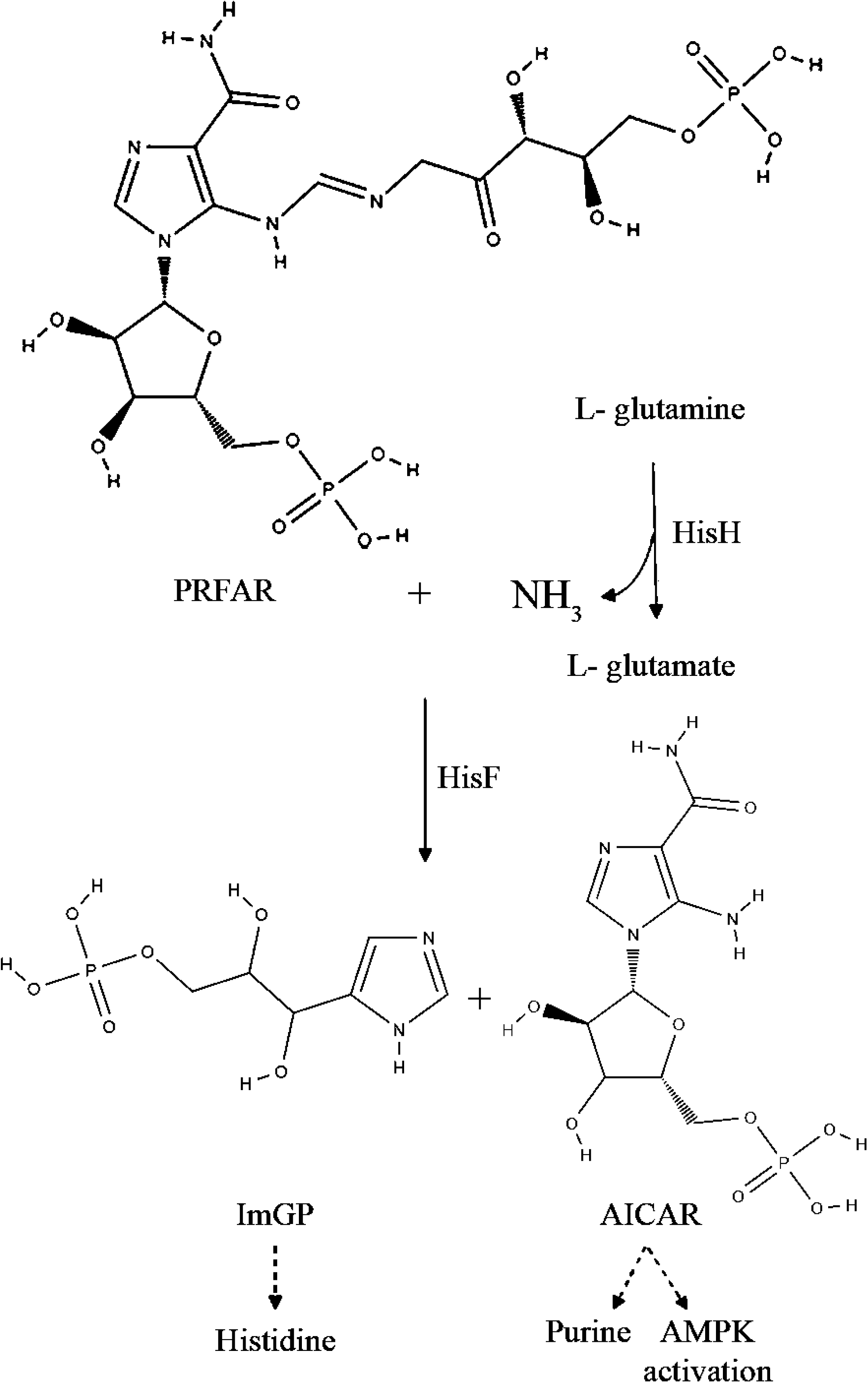
Reactions catalyzed by HisH and HisF. The ammonia molecule required for this reaction is provided by the glutaminase HisH which transfers nitrogen from L-glutamine to form L-glutamate. Later, PRFAR is converted by HisF into ImGP and AICAR. The second product of the reaction, AICAR, is further used in *de novo* purine biosynthesis and AMPK activation.

One of the products of HisF, AICAR, an analog of adenosine monophosphate (AMP), is capable of stimulating AMP-activated protein kinase (AMPK) activity. Both small molecules, AICAR monophosphate and AMP, trigger a conformational change in the AMPK complex that allows further activation by phosphorylation of Thr-172 (6). The central regulator of energy homeostasis, AMPK, is an enzyme that participates in the cellular response to metabolic stress, being considered as an important therapeutic target for controlling different human diseases (6).

Once activated, AMPK phosphorylates numerous metabolic enzymes causing a global inhibition of biosynthetic pathways and the activation of catabolic pathways, thus generating and conserving energy (7). It has been proven that AICAR, beyond the AMPK stimulation activity, is also able to inhibit the lipopolysaccharide-induced production of proinflammatory cytokines. The treatment with an adenosine kinase inhibitor was able to block the ability of AICAR to activate AMPK, preventing the inflammation inhibition in mice mesangial cells. (8, 9). Other authors have also described the role of AICAR in the regulation of inflammation (8, 10).

First, an experimental model of pneumonia in mice was employed to describe the transcriptome of the *A. baumannii* ATCC 17978 strain during the course of the infection, as previously published (11). A bronchoalveolar lavage (BAL) was performed to obtain bacteria suitable for RNA extraction (*in vivo* samples). RNA extracted from bacteria grown in LB medium was used as experimental control (*in vitro* samples). All experiments were done in accordance with regulatory guidelines and standards set by the Animal Ethics Committee (CHUAC, Spain, project code P82). Total RNA was used for RNAseq analysis (Illumina, Biogune, Spain). Raw data were deposited in the GEO database under the accession code GSE100552. Gene expression profiles were determined and analyzed as previously described (11). Transcriptomic analysis revealed that the A1S_3245 gene was over-expressed in bacteria isolated during the lung infection (7.2-fold change), compared with bacteria grown *in vitro*. Therefore, the aim of the present work was to study the role of this gene in the pathogenesis of *A. baumannii.*

Thus, the isogenic deletion derivative strain Δ3245 was obtained from the *A. baumannii* ATCC 17978 strain by double crossover recombination using the suicide vector pMo130, as described previously (11, 12). The upstream and downstream regions flanking the A1S_3245 gene were PCR-amplified and cloned into the vector pMo130 using the primers shown in Table S1.

Phenotypes of the parental and the mutant Δ3245 strains were compared through several *in vitro* assays previously described (11, 13), these including determination of biofilm, attachment to A549 human alveolar epithelial cells and motility abilities, as well as analysis of fitness and antimicrobial susceptibility (by disk diffusion assays), and no significant differences were observed (data not shown). Also, survival rates of A549 alveolar epithelial cells infected with the ATCC 17978 and the mutant Δ3245 strains were analyzed finding no differences.

A huge epithelial surface in contact with the inspired air makes lungs particularly susceptible to infection. This implies that respiratory tract must present wide defense mechanisms, such as the anatomical barriers of the nose or the phagocytic cells of alveoli. The cytokine IL-6 is involved in the regulation of inflammatory responses during bacterial infection and high IL-6 concentrations are detected in BAL fluids of patients with pneumonia (14). In murine models of pneumonia, IL-6 plays an important role in antibacterial host defense and in the regulation of the cytokine network in the lung (15). Thus, acute pulmonary inflammatory response caused by local exposure to bacterial lipopolysaccharide is regulated by inflammatory mediators such as IL-6.

Therefore, immunoassays were done to detect the cytokine IL-6 in macrophages RAW 264.7 infected with the parental and the mutant strain (MOI of 350), analyzing cell supernatants at 2, 6 and 20 h post-infection (N=5). IL-6 was measured by ELISA (16) using the Murine IL-6 ELISA Kit (Diaclone, France). Data indicated that the mutant strain produced more IL-6 than the parental strain (*p* = 0.01) at 2 h post-infection, disappearing this effect after 6 h post-infection, when no significant differences were found (Figure 2 A).

**Figure 2.**
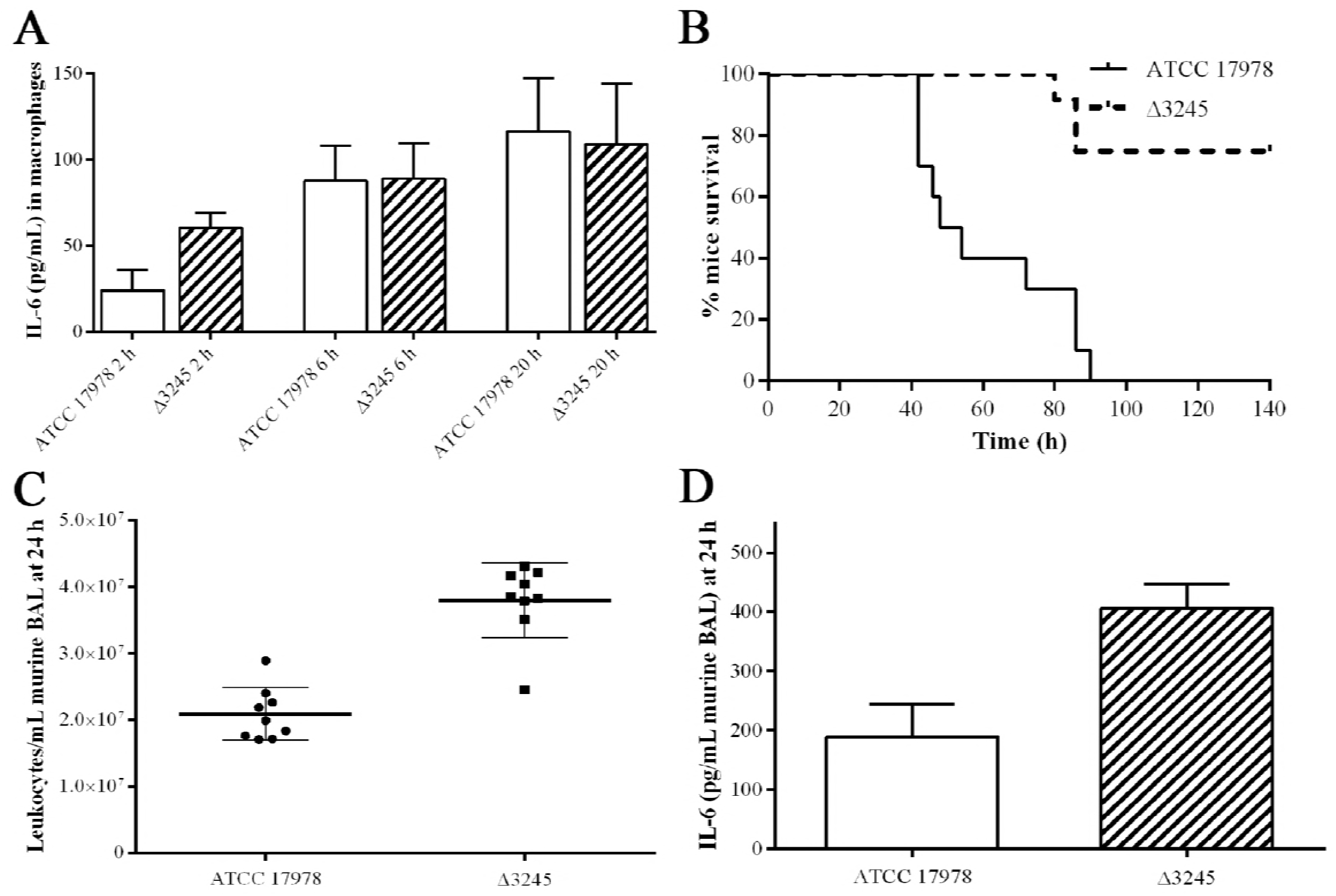
*In vitro* and *in vivo* assays using the parental ATCC 17978 and the mutant Δ3245 *A. baumannii* strains. A) Amount of IL-6 at 2, 6, and 20 h post-infection in the cell-free supernatant of macrophages RAW 264.7 (N=5). B) Survival rates in a murine pneumonia model (N=10). C) Total leukocytes counts in BAL from mice infected lungs (N= 7). D) Amount of IL-6 at 24 h in BAL from mice infected lungs (N=7).

Furthermore, a pneumonia model using BALB/c mice (N=10), infected by intratracheal inoculation with 40 µL of bacterial suspensions of 3 × 10^9^ CFU/mL of sterile saline solution and 10% porcine mucin (wt/vol) (Sigma-Aldrich) mixed at 1:1 ratio, was also used to analyze the role of the A1S_3245 gene in virulence. Data showed that mice infected with the mutant strain presented a significant greater survival rate than mice infected with the parental strain (*p* > 0.0001) (Figure 2 B).

BALs were performed in a murine pneumonia, 4 and 24 h after the challenge, to determine the total leukocyte cell counts (N=7). Cells were fixed and stained with Diff-Quick Stain (Thermo-Scientific, USA). Counts were performed using a microscope (Olympus, Japan) and the software Cell Sens Dimension (Olympus). Results showed *ca.* double counts of leukocytes from those lungs of mice infected with the mutant strain than in those infected with the ATCC 17978 strain (*p* > 0.001) at 24 h (Figure 2 C). Murine BALs obtained at an early stage post-infection did not show differences in the amount of leukocyte counts (data not shown).

Other aliquots of the above mentioned BALs (N=7) were centrifuged 1000 × *g* for 10 min and the cell-free supernatants were stored at −20°C until use. Supernatants were used to measure IL-6 and revealed that the IL-6 concentration was higher in BALs from mice infected with the mutant than in those infected with the parental strain (*p* = 0.007) at 24 h post-infection (Figure 2 D). In contrast to the infection caused on macrophages, no significant differences were observed at the early stage post-infection in mice. This fact, as expected, reflects that the immune system takes longer to reoccupy and express cytokines in mice than in the case of a direct infection on macrophages.

In addition, a murine sepsis model (N=10), where mice were inoculated with 100 µL of bacterial suspensions containing 75 × 10^7^ CFU/mL, was performed as previously described (17). Also, a *Galleria mellonella* infection model (N=10) was also carried out, where caterpillars (Bio Systems Technology, UK) were infected with 2 × 10^4^ CFU/larvae and virulence of the strains was evaluated analyzing the survival time, as described before (11). No changes were observed when the murine sepsis or the *G. mellonella* infection models were performed (data not shown).

Student’s *t*-test was performed to evaluate the statistical significance of the observed differences in all assays, except in the survival assays, where the survival curves were plotted using the Kaplan-Meier method and analyzed using the log-rank test. The *p* values ≤ 0.05 were considered statistically significant. All assays were performed at least by triplicate.

The *hisF* gene from *A. baumannii* ATCC 17978, found as over-expressed during the course of a pneumonia infection, is involved in virulence, which places it as a new potential target for antimicrobial therapies. Taking into account the data obtained here, the expression of the *hisF* gene seems to decrease the innate immunity and the inflammatory responses, which could partly explained the persistence ability of the strain in the lung.

## TRANSPARENCY DECLARATIONS

None to declare.

